# Synaptic Competition Enhances the Discrimination of Interfering Memories

**DOI:** 10.1101/2022.09.15.508061

**Authors:** Chi Chung Alan Fung, Tomoki Fukai

## Abstract

Long-term plasticity is the primary mode of synaptic weight updating. However, other updating modes include modification of synaptic connections across neurons. The synaptic competition found during adult neurogenesis is an example. In this process, adult-born neurons are integrated into the existing neuronal pool by competing with mature neurons for synaptic connections from the entorhinal cortex. Evidence suggests that adult neurogenesis is critical for discriminating similar memories, i.e., memories with considerable interferences. This computational study aims to link synaptic competition with pattern separation. We show that synaptic competition and neuronal maturation are crucial for separating considerably overlapping memory patterns. Furthermore, we demonstrate that a feed-forward neural network trained by a competition-based learning rule can outperform a multi-layer perceptron trained by the backpropagation algorithm, especially when the sample size is small. These results unveil the functional implications and potential applications of synaptic competition in neural computation.

## Introduction

Long-term synaptic plasticity such as spike-timing-dependent plasticity (STDP) ***Markram et al***. (***1997***); ***qiang Bi and ming Poo*** (***1998***); ***Sjöström et al***. (***2001***) have been considered to play a vital role in memory formation ***Abraham et al***. (***2019***). STDP suggests how a neuronal network could learn the appropriate connection strengths from temporal correlations between presynaptic and postsynaptic neuron pairs. A simplified picture proposed by Hebb suggests that neurons with positively correlated activities should be connected more strongly and vice versa ***Hebb*** (***1949***). This rule is known to be the Hebbian learning rule. Although the Hebbian rule is a good summary of long-term synaptic plasticity, the entire picture of the learning process in neuronal networks has yet to be completed ***Keck and Josselyn*** (***2021***).

Synaptic competition is another form of synaptic updating rules that may cause long-term changes in neuronal connections. Recent evidence suggests that synaptic competition occurs during adult neurogenesis in the hippocampus ***Adlaf et al***. (***2017***); ***Miller and Sahay*** (***2019***). The ablation of adult neurogenesis may impair some cognitive functions and cause mental disorders ***Anacker and Hen*** (***2017***), suggesting its importance in supporting normal brain functions. Although dendritic competition for afferent inputs has been known in the retina during normal development ***Perry and Linden*** (***1982***), why neurons compete for synaptic projections is unlikely to be limited within the normal development of neuronal circuits. Synaptic competition for pre-synaptic input likely plays a crucial role in neural information processing in the brain.

The birth of new neurons was previously considered never to occur during adulthood. Adult neurogenesis has been reported in the dentate gyrus (DG) ***Gould et al***. (***1990***); ***Kuhn et al***. (***1996***); ***Kempermann et al***. (***1997***); ***van Praag et al***. (***1999***), which is an entry terminal of the hippocampal formation, and the olfactory bulb ***Carleton et al***. (***2003***); ***Curtis et al***. (***2007***). DG receives excitatory synaptic input from the entorhinal cortex ***Witter*** (***2007***) and projects excitatory output to the hippocampal area CA3 ***Senzai*** (***2019***), and hence is a vital relay in the trisynaptic circuit ***Andersen*** (***1975***). Various factors that regulate the birth rate of new neurons have been shown in the adult hippocampus ***Gould et al***. (***1990***); ***Kuhn et al***. (***1996***); ***Kempermann et al***. (***1997***); ***van Praag et al***. (***1999***). Although its functional role has yet to be fully clarified, adult neurogenesis is essential for behavioral tasks that require the discrimination of similar memories ***Miller and Sahay*** (***2019***). Mice with ablated adult hippocampal neurogenesis fail to discriminate memories with significant mutual interference ***Clelland et al***. (***2009***); ***Pan et al***. (***2012***); ***Zhang et al***. (***2014***); ***Zhuo et al***. (***2016***). Similar effects were observed across different approaches in the ablation or the suppression of adult neurogenesis ***Pan et al***. (***2012***); ***Wojtowicz et al***. (***2008***); ***Burghardt et al***. (***2012***); ***Garthe et al***. (***2009***); ***Swan et al***. (***2014***). Furthermore, adult neurogenesis may reduce anxiety-like behaviors ***Anacker and Hen*** (***2017***), which was shown by using pharmacologically diverse antidepressants to promote adult neurogenesis in rodents ***Surget et al***. (***2008***); ***Malberg et al***. (***2000***); ***Banasr et al***. (***2006***); ***Dagytė et al***. (***2010***), non-human primates ***Perera et al***. (***2011***), human post-mortem brain tissue ***Boldrini et al***. (***2009***, 2012), and human hippocampal progenitor cells *in vitro* ***Anacker et al***. (***2011***).

Adult neurogenesis has not been a topic of extensive computational studies, but several computational models already exist in the literature. Some computational models focused on the role of neuronal turnover in memory processing rather than that of synaptic plasticity and synaptic competition ***DeCostanzo et al***. (***2019***). The model suggested that neuronal turnover significantly reduces the dimension of memory representations in the neural network of DG. A network model combining neuronal turnover and Hebbian learning was proposed, but this model did not also address the computational implications of synaptic competition ***Becker*** (***2005***). Another modeling study considered most biological features of adult neurogenesis ***Aimone et al***. (***2009***). However, the computational advantage of synaptic competition was not explicitly examined and remained elusive.

This study focuses on clarifying whether and how synaptic competition improves pattern separation. We begin by comparing performance in pattern separation between various learning rules, including synaptic competition and the conventional weight-updating rules of long-term potentiation (LTP) and long-term depression (LTD). We consider toy problems with overlapping input patterns to show the properties of different learning rules, and propose a learning algorithm that implements synaptic competition. Then, we train a multi-layer network adopting this learning rule on a real-world task, pattern separation on the MNIST dataset, and demonstrate that this network model can outperform a multi-layer perceptron trained by the back-propagation algorithm ***Rumelhart et al***. (***1986***). Notably, the synaptic competition-based learning rule works exceptionally well when only a small number of samples is available for training. Thus, our results suggest the advantage of synaptic competition over conventional Hebbian learning in memory formation with small samples or small trial numbers.

## Results

### LTP alone over-generalizes overlapping patterns

To begin with, we study a toy model to clarify how long-term potentiation modifies synaptic weights for overlapped patterns. As illustrated in Fig. 1(A), we model the long-term potentiation by the Hebbian learning rule: when pre-synaptic and post-synaptic neurons are coactivated, synapses between them should be strengthened. We will describe the mathematical details of the learning rule later. Each sample pattern *ξ* for training is generated as the sum of a pattern selected randomly from four fundamental patterns (Fig. 1(D)) and a noise vector (see Methods). These fundamental overlapped patterns are mathematically given as

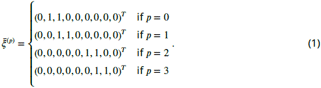

**Figure 1.**
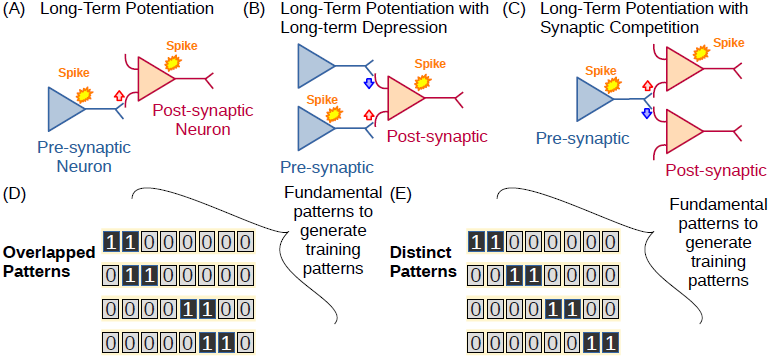
Synaptic plasticity and synaptic competition. (A) LTP is illustrated schematically. (B) Combinations of LTP and LTD occur on a single neuron. (C) LTP is combined with synaptic competition in which synapses compete for presynaptic resources.

The parameter η controls the magnitude of noise. Note that patterns generated from pattern *p* = 0 overlap with those from pattern *p* = 1 (group 1), while patterns obtained from pattern *p* = 2 overlap with those from pattern *p* = 3 (group 2). There is no overlaps between patterns belonging to the different groups.

To implement the Hebbian learning model, we consider a two-layer neural network in which the first layer is an input terminal and the second layer is an output terminal (see Methods). Activity of the *j*-th input unit is denoted by *x*_*j*_ while that of the *i*-th output unit by *y*_*i*_, and the weight of synapse connecting the output unit *i* and input unit *j* is denoted by *w*_*ij*_. During learning, the synaptic weights evolve as

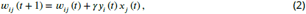

where *t* specifies the time of presentation of the *t*-th sample pattern and *γ* = 0.1 is the learning rate. The values of *w*_*ij*_ are randomly initialized. In the training process, sample patterns are presented one by one. We normalize input synaptic weights on each output unit if the maximum weight exceeds 1 on the output unit:

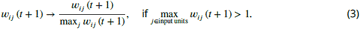

In this setting, we can easily deduce the trained weights; that is, changes in input synaptic weights averaged over time should be proportional to the averaged input patterns:

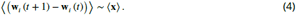

This implies that the presentation of an input pattern rotates the synaptic weight vector in the direction of the input pattern. Thus, for the patterns given in Eq. (1), the resultant weights will be given as

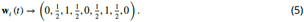

As the number of trained samples increases, input weights onto output units will become similar to each other. The equilibrium weights will show no difference between the two fundamental patterns in Eq. (1) and mutual information between the output **y** and input **x** will also vanish. Fig. 2(A) shows typical examples of input synaptic weights obtained for a representative neuron by numerical simulations. The numerical results agree well with those predicted in Eq.(5) except for slight deviations from the expected symmetric shape. The deviations are due to the sequential training of the weights. Neurons favor the patterns that appear earlier in the training sequence. We note that input synaptic weights on other neurons are almost identical to those shown in Fig. 2(A). Actually, the means of synaptic weights are more or less identical (Appendix 1-Fig. 1(A)) and their standard deviation across neurons are almost zero (Appendix 1-Fig. 1(B)).

**Figure 2.**
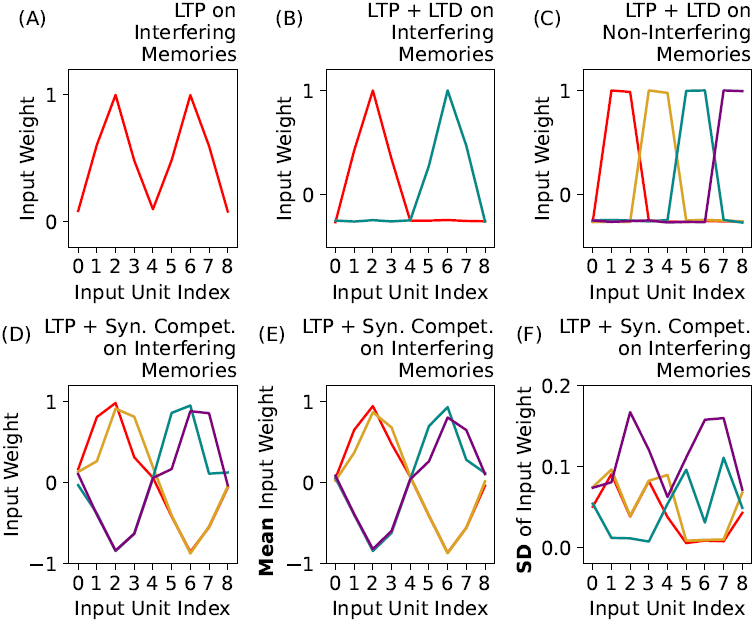
The profiles of input connection weights trained by different training rules under different scenarios. (A) A representative profile trained by LTP with samples derived from the overlapping fundamental patterns given in Eq. (1). (B) Representative profiles trained by LTP and LTD with overlapping fundamental patterns. Two weight profiles peaking at input units 2 (red) and 6 (cyan) are obtained. (C) Representative profiles trained by LTP and LTD with samples of the non-overlapping fundamental patterns given in Eq. (8). Four distinct profiles with different peak locations appear. (D) Representative profiles trained by LTP and synaptic competition with samples of the overlapping fundamental patterns. The profiles are categorized into two groups (red and orange vs. purple and cyan) according to the grouping of excited and inhibited input units. (E, F) The mean and standard deviation of weight profiles over output units for the simulations shown in (D), respectively.

### LTP and LTD jointly separate non-overlapping patterns

In the previous example, the LTP-like Hebbian learning rule tends to over-generalize the training patterns. With the input synaptic weights plotted in Fig. 2(A), the responses of output neurons would not distinguish the different fundamental patterns. In this sub-section, we incorporate LTD to examine if combining LTP and LTD in learning rules improves the resolution of separation between overlapped patterns. Figure 1(B) schematically illustrates the LTD used in this study.

To this end, we modify the learning rule in Eq. (2) as follows:

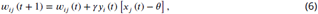

where the additional term *θ* controls the intensity of LTD. Input synaptic weights undergo LTD when the corresponding presynaptic activities are weaker than the threshold value. Equation (6) has a similar structure to Eq. (2), and the average change of input weights would be

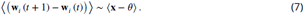

The above equation is similar to Eq. (4) except that inhibitory connections can emerge due to the *θ* term. Now, input patterns corresponding to *p* = 0 and 1 and those corresponding to *p* = 2 and 3 excite non-overlapping input-unit subgroups. Therefore, Eq. (7) implies that input weights and output neurons will evolve into two subgroups, one responding selectively to the former grouped patterns and one responding selectively to the latter grouped patterns (Fig. 2(B)).

The results suggest that LTD helps output neurons to distinguish the distinct groups of input patterns (i.e., the group for *p* = 0, 1 and that for *p* = 2, 3). From another perspective, we may say that LTD generalizes different but similar patterns (i.e., patterns for *p* = 1 and those for *p* = 2, etc.) into the same category. The means and standard deviations of input weights of the different groups are shown in Appendix 1-Figs. 1(C) and 1(D), respectively. Input weights take identical values on neurons tuned to the same group of overlapping patterns.

The above results showed that LTD can separate non-overlapping groups of patterns. To further verify the implications of LTD in pattern separation, we investigated how the pattern separation occurs when the fundamental patterns are not mutually overlapped (Fig. 1(E)):

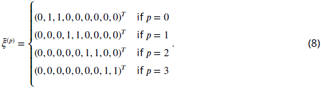

Input weights are trained by Eq. (6). Since the sample patterns corresponding to *p* = 0, 1, 2 and 3 have no overlaps and they are presented in a random sequence, output nodes inhibited by a sample pattern can be excited by some other patterns. As a result, input weights separate into four groups:

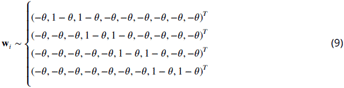

Input weights resultant from the simulation are shown in in Figs. 2(C) and Appendix 1-1(E-F). As suggested by Eq. (9), the numerical results yielded four types of input weights. These analytical and theoretical results revealed that the combination of LTP and LTD can organize input weights to distinguish non-overlapping patterns. However, as shown previously, this learning rule generalizes but does not distinguishes overlapping patterns.

Before we study synaptic competition, we analyze the network model trained on non-overlapping input patterns given in Eq. (8) with the LTP-based learning rule in Eq. (2). In this case, input weights should self-organize as

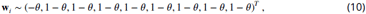

which implies that LTP-based rule classifies everything into a single category. Namely, pattern separation fails. This result tells us that LTD should counter-balance the over-generalization by LTP in pattern separation.

### Synaptic competition differentiates overlapping patterns

In the previous subsections, we demonstrated that learning rules combining LTP and LTD tend to over-generalize similar input patterns and cannot differentiate overlapping input patterns such as given in Eq. (1). As input patterns generally have overlaps in many real-world problems, this result severely limits the validity of the feedforward network model in pattern separation. Another type of weight updating rule is necessary to differentiate mutually interfering patterns.

The extreme over-generalization may occur when different input patterns do not compete with each other for getting resources for learning, that is, synaptic connections. Synaptic learning rules without competition are also far from realistic as they assume that infinite resources are available at each synapse. In reality, there should be an upper bound of the number and/or total strength of synaptic connections.

We searched for an efficient mechanism of synaptic competition by modeling some properties of adult neurogenesis, which occurs in the dentate gyrus and olfactory bulb. Adult neurogenesis is considered crucial for pattern separation in episodic memory and odor discrimination. In those brain regions, the maturation of newly born neurons is an important process for developing synaptic connections from neurons in input layers. During this process, newly born neurons compete for input synaptic connections with existing neurons. Furthermore, the expansion of synaptic contacts terminates on matured newborn neurons. Below, we implement these two properties in the toy model.

Considering that the shortage of resources would affect the growth of synapses, we may model the maturation of synaptic contacts by terminating LTP if the sum of input weights on each neuron exceeds a certain threshold. We can describe the maturation process by the following equation:

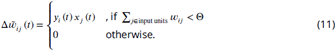

This equation models the degradation of LTP after maturation ***Schmidt-Hieber et al***. (***2004***). We model synaptic competition for pre-synaptic resources as

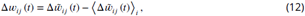

in terms of the long-term average of weight changes on each neuron. This equation should not be confused with homeostatic plasticity regulated by the postsynaptic activity ***Kaleb et al***. (***2021***). The updating rule for *w*_*ij*_ is given as

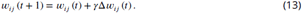

Then, we renormalize all synaptic weights from input unit *j* if the largest weight from the unit exceeds unity:

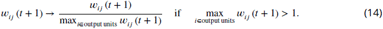

Unlike Eq. (3), Eq. (14) normalizes the net output of input units.

While eq. (11) integrates all input weights that can excite an input unit, eq. (12) induces competition on a postsynaptic neuron between input weights from different input neurons. The competition breaks the symmetric configuration of input weights in Fig. 2(A) during training. Representative input weights are shown in Fig. 2(D), which reveals four types of tuning curves. Two of them have peaks at the input unit 2 and another pair at the input unit 6. This configuration, with peaks at the units 2 and 6, is similar to that in Figure 2(B). However, the competition between input weights segregates input units, inducing a broader tuning pattern in the output neuron population. Consequently, we can now categorize input weights in Figure 2(D) by their locations of the second-highest values. The means and standard deviations of the input weights for different categories are shown in Figs. 2(E) and 2(F), respectively. These results have important implications for pattern separation as they suggest synaptic competition, unlike LTD, enables the network model to differentiate similar input patterns. Below, we will further compare the effects of synaptic competition and those of LTD in a more comprehensive fashion.

### Principle component analysis for synaptic competition

For a clear comparison of input weights between the different algorithms, we employed principal-component analysis (PCA) to reduce the dimension of input weights. Here, we used the standard PCA without whitening and show the resultant principal components (PCs) corresponding to the three largest eigenvalues. Figures 3(A) and 3(B) show the projections of input weights on the first and second principal components or on the second and third principal components, respectively. We labeled the projected input weights according to their locations of the second-largest synaptic weight. The input weights projected onto the first principal compartment (PC1_compet_) clearly separated the input weights corresponding to patterns *p* = 0 and 1 from those corresponding to patterns *p* = 2 and 3. Furthermore, the input weights tuned to patterns *p* = 0 and *p* = 1 are well separated in the subspace spanned by the second and third principle components, (PC2_compet_) and (PC3_compet_). Similarly, input weights tuned to patterns *p* = 2 and *p* = 3 were segregated in the same subspace. Figure 3(C) shows the same plots in a 3D space to visualize the clear separation of the four groups of input weights. These results demonstrate how the input weights trained through the combination of LTP and synaptic competition distinguish similar patterns. In contrast, the input weights trained by LTP and LTD without synaptic competition are only tuned to two groups separating non-overlapping patterns (Fig. 3(D)). The minor differences between overlapping patterns are largely ignored. Altogether, our results clarify the crucial role of synaptic competition in separating similar patterns.

**Figure 3.**
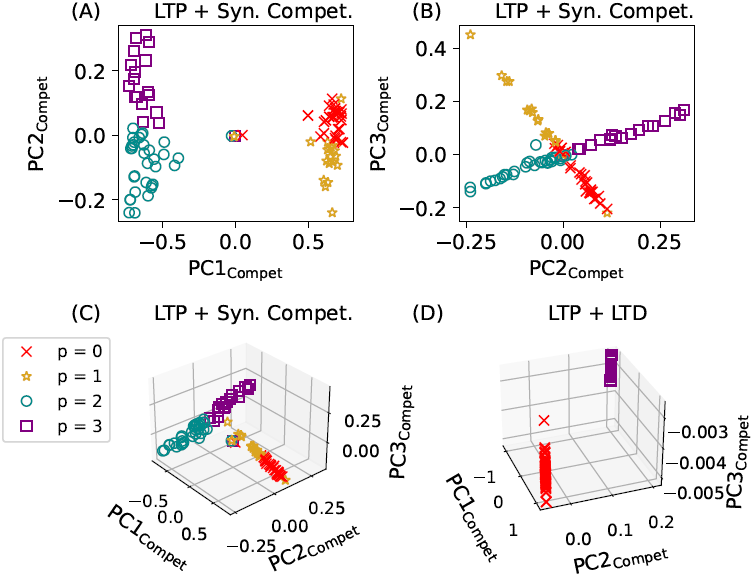
Input weights in the toy model trained by synaptic competition. Pseudocolor code indicates the preferred fundamental patterns defined by the positions of the second-largest input weights on individual output neurons. (A) The trained input weights are projected on the first two leading PCs, i.e., PC1_Compet_ and PC2_Compet_. (B) The trained input weights projected on PC2_Compet_ and PC3_Compet_. (C) A three-dimensional plot is shown for the trained input weights in (A) and (B). (D) Input weights trained by LTP and LTD are plotted in a 3D space spanned by the leading PCs obtained for synaptic competition.

To further confirm the unique role of synaptic competition in pattern separation, we conducted PCA on output neuronal activities, *y*_*i*_, obtained for patterns generated from the fundamental patterns. When the network model is trained by LTP and synaptic competition, the projected neuronal activities corresponding to different (both overlapping and non-overlapping patterns) fundamental patterns are clearly distinguished (Fig. 4(A)). However, if the network model is trained by LTP and LTD without synaptic competition, they can only separate non-overlapping patterns but not overlapping patterns (Fig. 4(B)). Output activities are significantly overlapped for overlapping fundamental patterns. We also show neuronal activities in a network trained only with LTP. In this model, projected neuronal activities generally fall into a single cluster, implying that such a network has little capability of pattern separation (Fig. 4(C)). The results shown here further confirmed that synaptic competition greatly enhances pattern separation in the toy model.

**Figure 4.**
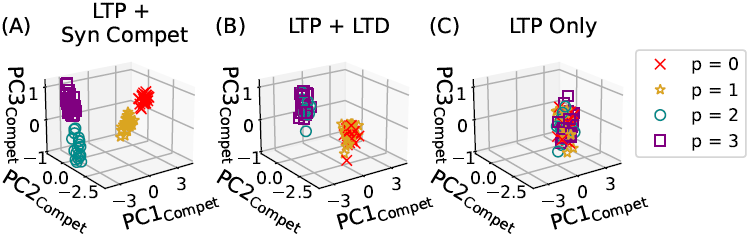
Neuronal activities in the toy model trained by synaptic competition. Pseudocolor code indicates the preferred fundamental patterns of output neurons. (A) The outputs of neurons encoding different preferred fundamental patterns are plotted in the 3D space of the leading PCs. (B, C) Similar plots are shown for networks trained by (B) a combination of LTP and LTD and (C) LTP alone.

### Distinct roles of maturation and competition in pattern separation

The synaptic competition rule proposed above consists of two components: maturation, i.e., Eq. (11), and competition for pre-synaptic resources, i.e., Eq. (12). One may ask which component is more important to achieve pattern separation shown in Fig. 4. To answer this question, we modify the learning rule defined by Eqs. (11) - (13). To examine if the maturation is important, we replace Eq. (11) with the simplest Hebbian rule defined in Eq. (2). Similarly, to check if the competition is important, we drop the second term in the right-hand side of Eq. (12). we trained the neuronal networks that underwent either of these modifications.

From the results shown in Fig. 5(A) and (B), we find that neither of the network models can perform pattern separation at a satisfactory level: neuronal activities projected on the leading principal-components exhibit a single cluster in each scenario. These results demonstrate that both maturation and competition on pre-synaptic resources are indispensable for pattern separation in the present simple task setting.

**Figure 5.**
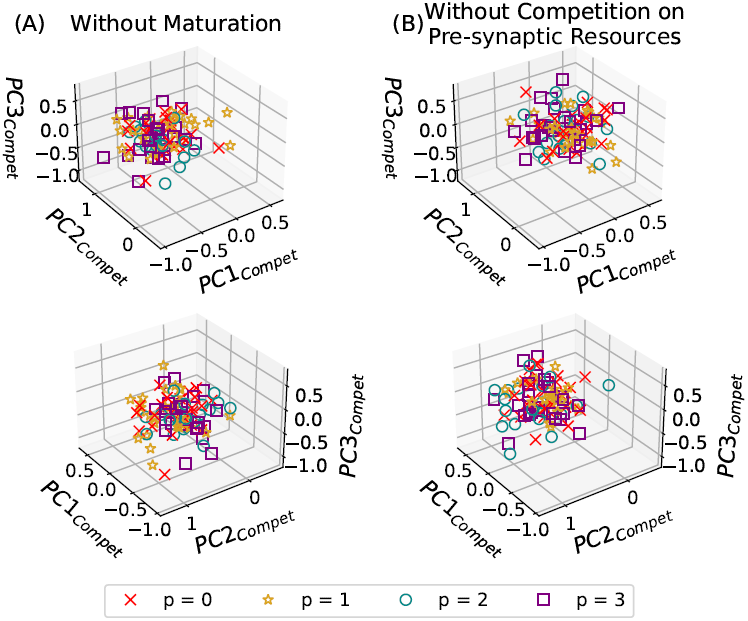
Neuronal activities in the toy model trained by disrupted synaptic competition rules. Pseudocolor code indicates the preferred fundamental patterns of output neurons. (A) Projections of neuronal activities on the PC space in a toy model trained by synaptic competition without the maturation term. Upper and lower panels show the same plot from different angles. (B) Neuronal activities are projected on the PC space in a toy model trained by synaptic competition without competition for pre-synaptic resources.

### Synaptic competition facilitates pattern separation with small sample sizes

The toy model demonstrated the unique role of synaptic competition in discriminating interfering memory patterns. However, the toy model trained to discriminate only four patterns is too simple to claim the validity of synaptic competition for pattern separation. One may ask whether the competition-based learning rule works similarly well on real-world classification tasks with more complex input patterns. In the following, we train a feedforward network model by synaptic competition to examine whether the network can perform the classification of hand-written digits.

To this end, we consider three-layer feedforward neuronal networks equipped with synaptic competition and the MNIST database of hand-written digits (from 0 to 9). We also consider a multilayer perceptron (MLP) for comparison in performance. The outline of our comparative study is explained in Fig. 6(A). Both our feed-forward neuronal networks for synaptic competition and MLP have 28^2^ units in their input layers and ten units in their output layer to discriminate the ten digits. Thus, the output layer supports one-hot coding, which gives a winner-take-all binary representation to each digit. In both models, neurons are rectified linear units (ReLU), that is, the activation function of neurons is a rectified linear function. The number of neuronal units in the middle layer is a parameter crucial for the comparison between the models. The differences between the two networks are the learning rules to train input and output weights. The weights are updated in the MLP network by the error-backpropagation algorithm (see Method). In the network for synaptic competition, input weights were updated by synaptic competition, while the least-square fitting determined output weights after the training of input weights (see Method).

**Figure 6.**
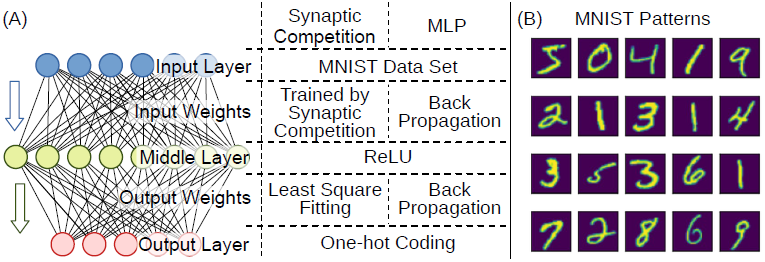
Performance test on a real-world classification task. (A) The structures of three-layer neuronal networks are compared. A competition-based model was trained by synaptic competition and least square fitting, while a multi-layer perception (MLP) was trained by the back-propagation algorithm. (B) Samples of hand-written digits in the MNIST data set.

We found that the synaptic competition-based learning rule boosts classification performance, especially when the sample size is small. There are in total 60,000 hand-written digits in the MNIST dataset. Some examples are shown in Fig. 6(B). In this work, we trained the network models on relatively small subsets of the dataset with 5,000, 10,000, and 50,000 samples. Machine learning often requires big data, which severely limits the applicability of such algorithms. Therefore, we are particularly interested in the situation where only a small number of samples are available for learning. Furthermore, each input pattern is exposed only once to the network model with synaptic competition but five times in total to the MLP network.Under this condition, which is more stringent for the competition-based network model than for the MLP network, we trained both network models with different sizes of the middle layer. To our surprise, on the smallest sample data, the network with synaptic competition always outperforms the MLP network for all sizes of the middle layer (5,000 samples, Fig. 7(A). When the sample size is doubled, the competition-based network model remains superior if it has a sufficiently large middle layer (10,000 samples, Fig. 7(B)). The minimal size of the middle layer further increases as the sample size is increased, but the competition-based network model still outperforms the MLP network for a large enough middle layer (50,000 samples, Fig. 7(C)). These results suggest the advantage of synaptic competition-based learning rules in real-world pattern classification tasks.

**Figure 7.**
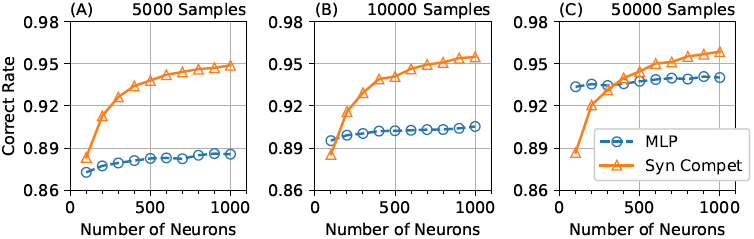
Correct rates of the competition-based model and MLP. The two networks were trained with (A) 5000, (B) 10000, and (C) 50000 samples from the MNIST dataset.

We demonstrate how synaptic competition provides an improved cue for classifying hand-written digits of the MNIST dataset. In the synaptic competition-based model, the tuning profiles of input weights on individual middle-layer neurons show interesting features. Figure 8 shows examples of the tuning profiles, which look like “blurred” digits. Unlike other classification algorithms, the digits the neurons have learned are not obvious from the tuning profiles. The configuration of excitatory connections likely captures only a tiny portion of each digit, implying that the trained neurons are tuned to some strokes of the digits. We further conducted PCA of neuronal activities in the network trained by synaptic competition. Figure 9(A) displays the input pattern distributions for digits 0 and 8 projected on the two-dimensional space spanned by their first two PCs. Similar distributions of PC1 and PC2 are shown in Fig. 9 (B) for middle-layer neuronal activities. While the distributions of the two digits are barely separated in the input layer, they are better separated in the middle layer. The projections of input patterns and middle-layer activities on the spaces of higher-order PCs suggest that some higher-order components also contribute to pattern classification (Appendix 1-Fig. 2). We show a more quantitative comparison across the different pairs of sample digits in Fig. 10, where we measured Mahalanobis distances between the clusters obtained for the two sample digits in different layers. In the lowest PC dimensions, digit pairs show significantly larger distances in the middle layer than in the input layer. In contrast, in the higher PC dimensions, distances between digit pairs are significantly smaller in the middle layer than in the input layer. The comparison results indicate that the synaptic competition-based learning rule concentrates the differences between different digits in lower PC dimensions. This property of synaptic competition improves the output layer’s classification ability.

**Figure 8.**
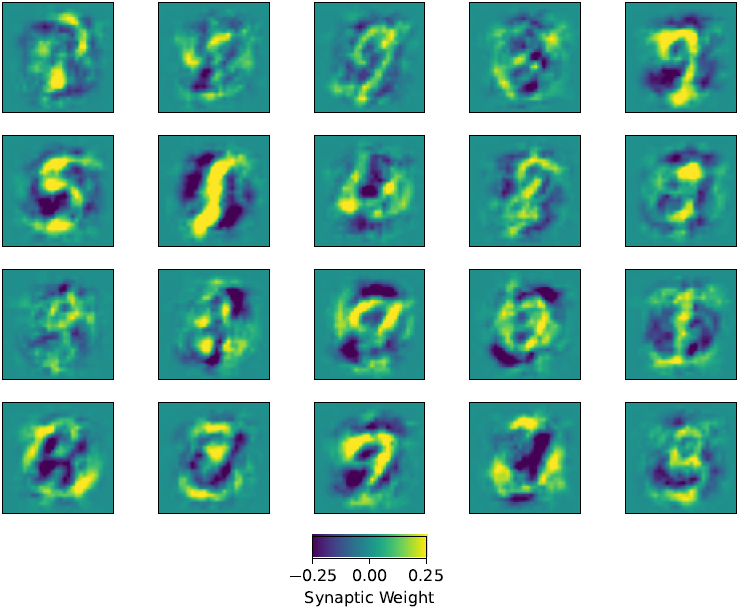
Neural representations in the competition-based network model. Examples of trained input weights on the middle layer are shown for the competition based network model in Fig. 6(A).

**Figure 9.**
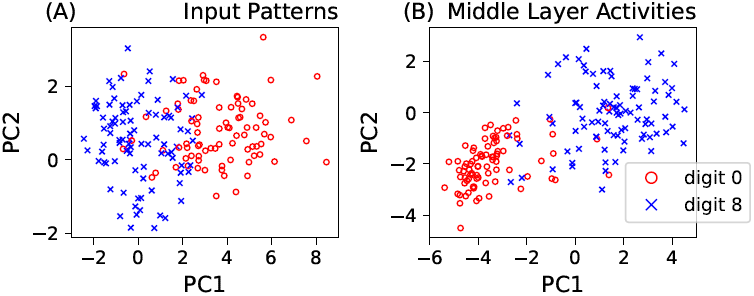
Representations of digits in the different layers of the competition-based network model. (A) PCs of input patterns are shown for hand-written digits 0 (red) and 8 (blue). (B) PCs of neuronal activities are shown for middle-layer neurons with preference to hand-written digits 0 and 8.

**Figure 10.**
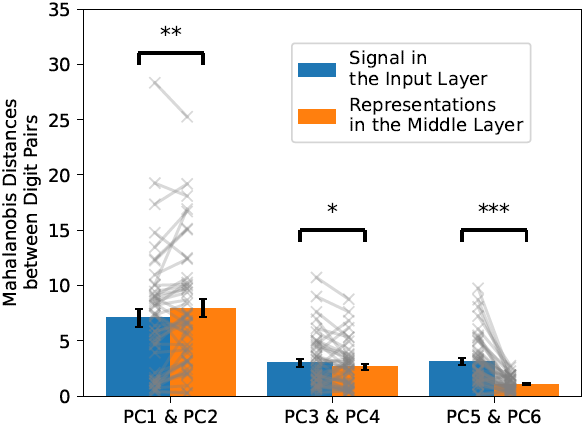
Separation of digit pairs in the competition-based network model. Mahalanobis distances between PCs of digit pairs are shown in the input layer (blue) and the middle layer (orange). The distances were calculated in the two-dimensional spaces spanned by “PC1 and PC2”, “PC3 and PC4”, and “PC5 and PC6”. Error bars show the standard deviations of means, and p-values were calculated by paired *t*-test: *** (*p* < 0.001); ** (*p* < 0.01); * (*p* < 0.05).

The layers trained by synaptic competition could provide extra separations for different classes.

Unlike the examples shown in Fig. 9, improvements in major PCs are less obvious for some combinations of digits, as shown by digit-by-digit PC comparisons in Appendix 1-Fig. 3. For most pairs of digits, the first two PCs of input patterns do not show an obviously improved separation compared with those obtained from the middle layer trained by synaptic competition. However, this is not necessarily negative news because the input patterns are somewhat separable before they undergo synaptic competition. To show this, we considered a network model without a hidden layer. Namely, we connected input units directly to output units and tuned the weights of direct connections by the least square fitting. This input-direct least square fitting resulted in 82.4% correct classification, compared with the correct rate of 91.5% for a network model with a 200-unit hidden layer trained by the synaptic competition rule. The synaptic competition rule outperformed the input-direct least square fitting for all digit classes (Fig. 11), implying that synaptic competition at the middle layer boosts separations between different digit classes. Furthermore, we compared classification performance between normalized random input weights and input weights trained by synaptic competition. We sampled the normalized random weights from a normal distribution with zero mean and unity variance. Synaptic competition outperformed random weight sampling in all digit classes, although the overall correct rate was as high as 85.8% for the latter. These results prove the advantage of synaptic competition as an algorithm doing pattern separation.

**Figure 11.**
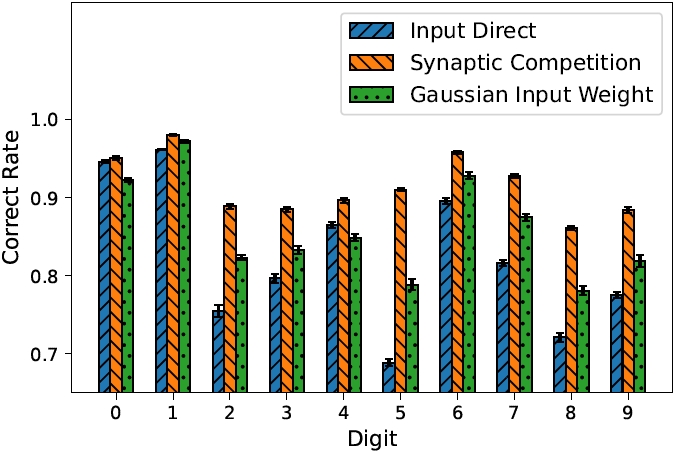
Performance comparison between the different algorithms. Separate comparison was made by using 5000 samples of each digit class. The number of units in the middle layer was 200 for synaptic competition and Gaussian input weight. Error bars indicate the standard deviations of means over 10 independent trials.

## Discussion

### Comparison with other models

The present work investigated how a feedforward network with synaptic competition for presynaptic resources separates interfering input patterns in an unsupervised way. In particular, our results suggest that synaptic competition greatly enhances pattern separation when the size of sample data is small. In addition, we exposed our network model to each sample of input patterns only once during training, aiming at mimicking biologically realistic situations of online learning. Although the biological details of synaptic competition are over-simplified, We formulated the core effects of our weight updating rule, namely, the maturation and competition terms, based on recent biological observations ***Miller and Sahay*** (***2019***). Therefore, we believe that our model captures the key properties of synaptic competition in biological neuronal networks.

Although learning by competition was reported in the literature, the model of synaptic competition shown in this study is novel in several aspects. A recent study proposed an algorithm in which different input patterns compete for projections to hidden layer neurons ***Krotov and Hopfield*** (***2019***). Although the algorithm worked well in numerical experiments with the MNIST data set, some input neurons developed projections to many hidden layer neurons, implying that many input patterns activate these input neurons. This overload to particular neurons makes such a model biologically less plausible. The model was also not tested on online learning with a small number of samples.

By contrast, the competition term in the present learning rule functions as a subtractive normalization term, suppressing the sum of outgoing weights of each input neuron. This prevents the overload of input neurons. Other models for synaptic competition mainly focused on competition for post-synaptic resources. For instance, in a model (e.g., ***Triesch et al***. (***2018***)), restrictions of resources in postsynaptic neurons may limit the sum of input weights on each hidden layer neuron. However, while the model shed light on the dynamics of physiological processes, it did not clarify the crucial role of synaptic competition in pattern separation.

Some numerical studies on adult neurogenesis explored the computational functions of neuronal turnover. We previously reported a network model in which input weights on newborn DG neurons were assigned randomly after their births ***DeCostanzo et al***. (***2019***). However, the model ignored long-term plasticity, which is known to play a non-negligible role in developing dendrites during the 4 to 6 weeks of the birth of new neurons. Another model considered long-term plasticity by using a Hebbian learning rule ***Becker*** (***2005***). However, the learning rule did not explicitly take the role of synaptic competition into account, missing the important characteristics of adult neurogenesis. Yet another model involved the concept of synaptic competition ***Aimone et al***. (***2009***), but did not reveal its nontrivial roles such as those studied here. Although our simplified model is far from a full spec model of neurogenesis, our numerical analyses revealed the essential contribution of synaptic competition to the discrimination of similar but different stimuli, a vital brain function. Furthermore, our model predicted that disabling synaptic competition impairs pattern separation only when stimuli are similar (see Fig. 4). This prediction is testable by future experiments.

### Potential application in machine learning

Neural networks play a pivotal role in machine learning and artificial intelligence. Investigations of neural pathways in the brain have often inspired novel ideas and techniques in machine learning. For example, the neocognitron invented by Fukushima was inspired by the anatomical organization and neural responses of visual pathway ***Fukushima*** (***1980***). Since then, a rich family of neural networks, namely, the convolutional neural network, has been proposed for the variety of tasks, ranging from classification ***Lecun et al***. (***1998***) to style transfer ***Gatys et al***. (***2016***). Recently, the coding of a modern machine learning algorithm for natural language processing was found to be consistent with experimentally observed brain signals ***Caucheteux and King*** (***2022***). These studies demonstrate how the discoveries and knowledge learned from the brain could advance methods in machine learning.

In the current study, neurons in the hidden layer compete for input signals and this competition is activity-dependent. This means that synaptic competition processes input activity patterns based on the nature of the patterns. Thus, synaptic competition is an unsupervised algorithm for training input weights of hidden units without specifying particular purposes. Further, the comparison between models shown in Fig. 7 reveals that synaptic competition significantly improves classification performance when the sample data size is small. We may intuitively understand this remarkable feature of presynaptically driven synaptic competition through its basic property. When synapses compete for input patterns, they will be more successful if they can discriminate minor differences between different input patterns. We speculate that this pressure enables our network model to discriminate mutually interfering input patterns. This ability of synaptic competition is quite useful for online training, which requires simultaneous learning and data collection. Furthermore, in Fig. 7, we obtained the results for synaptic competition just after one training cycle, whereas those for back-propagation required five cycles. These results suggest that the synaptic competition rule can generate efficient and effective coding of input patterns. Although the particular network architecture shown in Fig. 6(A) may not suit other tasks, the learning rule proposed in this study can widely be used.

In sum, we have demonstrated that synaptic competition can enhance pattern separation. It suppresses the effect of interference across similar input patterns. Our results suggest that synaptic competition makes similar patterns more distinguishable during the sensitive period of newly-born neurons in adult neurogenesis. We have also shown that synaptic competition gives a promising algorithm for unsupervised learning in real-world classification tasks. Our findings will contribute not only to advancing the understanding of the adult neurogenesis but also to developing novel online machine learning methods for difficult classification tasks.

## Methods

### The Neuronal Network and Training Patterns Generated from Fundamental Patterns

The network is defined by

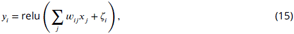

where *x*_*j*_ is the input value at input node *j, y*_*i*_ is the activation of node *i, w*_*ij*_ is the coupling between them and ζ_*i*_ *j* is the input noise.

The sample patterns were constructed from fundamental patterns:

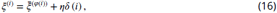

where 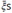 are fundamental patterns given in Eq. (1) or Eq. (8), *φ* is the random mapping from *i* to {0, 1, 2, 3}, *η* = 0.1 is the magnitude of the noise, and *δ* is a noise vector with entries between 0 and 1.

### Training Process with Different Training Rules

There are 1000 training samples were generated by Eq. (16) for training processes aimed to show the difference between training rules with long-term potentiation, long-term potentiation with long-term depression, and long-term potentiation with the synaptic competition.

In training, all input weights were initially zero. In each iteration, training patterns were inputted into the network in a one-by-one manner. After each input, the neuronal activity *y* will be updated. Then input weights were updated according to Δ*w* in the equations accordingly. In particular, input weights were trained in only one iteration.

The parameters used in various studies are presented in the following. For long-term synaptic potentiation, in Eq. (2), the learning rate, *γ*, was fixed to be 0.1. For long-term synaptic potentiation and long-term synaptic depression, in Eq. (6), the learning rate, *γ*, was also fixed to be 0.1, and the level of long-term depression, *θ*, was chosen to be 0.25. For long-term synaptic potentiation with synaptic competition, in Eq. (13), the learning rate, *γ*, was fixed to be 0.1. In Eq. (11), the threshold for maturation, Θ, was chosen to be 1.0.

### The Classification-task Comparison

#### The Data Set

The hand-written digit data set MNIST consists of 70000 digits. The size of the input is 28 × 28 = 784 pixels. The numbers of training samples were 5000, 10000, and 50000. The rest of the digits were reserved to be in the testing set. The corresponding labels of those digits were one-hot coding. One-hot means that the expected output of the feedforward network is an array having 10 binary entries and only one of the entries is one.

#### Neuronal Network Trained by Synaptic Competition

In the training of the input weights shown in Fig. 6(A), the training digits were inputted into the network one by one. After each input, the input synaptic weights will be updated by the synaptic competition rule. The parameters for the synaptic competition rule were *γ* = 0.01 and Θ = 1.0.

The output weights, **w**^out^, is determined by least-square fitting. Let **x**^training^ is a matrix whose columns are training input patterns, and **w** is the matrix for the trained input weights. We have**y**

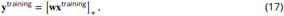

where [⋅]_+_ is an element-wise rectified linear function. By using the neuronal activity **y**^training^ and the expected one-hot coding **z**^training^, we can determine the output weights **w**^out^ as follow.

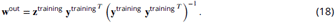

In the assessment of classification performance, digit patterns from the testing pool were inputted into the input layer of the network. The coding of the output layer will be simply the winner-take-all rule. The correct rate was calculated by the ratio of the number of successful predictions and the size of the testing pool.

#### Multi-layer Perceptrons Trained by Back-propagation

The multi-layer perceptron network used in this study was constructed by an interface, namely Keras ***Chollet et al***. (***2015***). The back-end behind Keras is Tensorflow. The middle layer of the network was constructed by a number of rectified linear units. The output layer is constructed by softmax functions.

In the training phase, the network was trained by the training routine included in Keras. The back-propagation was done by stochastic gradient descent (SGD) optimizer. The loss function was categorical cross-entropy. There were five training epochs on the whole training set before testing. During testing, the calculation was similar. The correct rate is calculated by the ratio between successful predictions and the size of the testing pool.

## Acknowledgments

This work is supported by KAKENHI nos. 19K16885 to CCAF, 18H05213 to TF, and 19H04994 to TF from the Japan Society for the Promotion of Science and by the start-up grant no.9610591 from City University of Hong Kong to CCAF.

## Appendix 1

### Supplementary Figures

**Appendix 1—figure 1.**
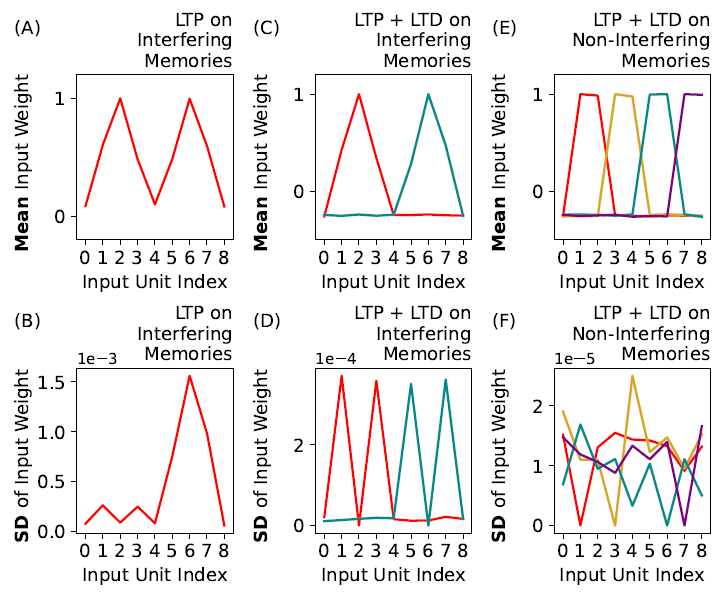
Means and standard deviations of input connection weight profiles trained by different long-term potentiation (LTP) and long-term potentiation with long-term depression (LTP + LTD) under different scenarios. (A) & (B): Mean and standard deviation (SD) of input weight profiles trained by long-term potentiation on overlapping patterns. (C) & (D): Mean and standard deviation (SD) of input weight profiles trained by long-term potentiation with long-term depression on overlapping patterns. (E) & (F): Mean and standard deviation (SD) of input weight profiles trained by long-term potentiation with long-term depression on distant patterns.

**Appendix 1—figure 2.**
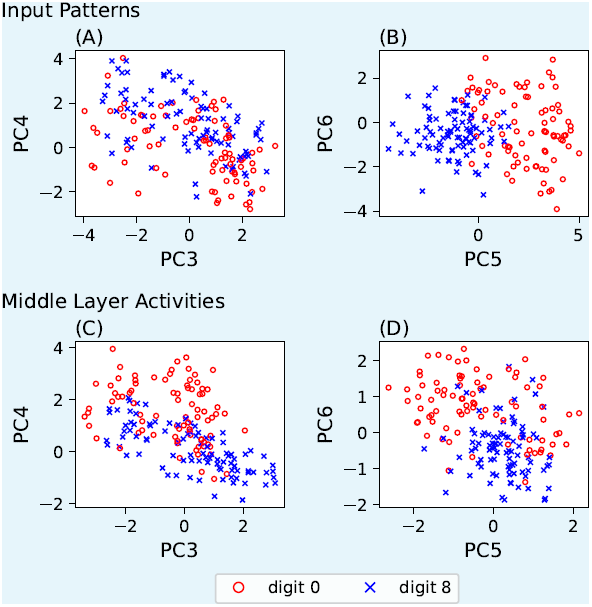
(A) & (B): Higher principal-component (PC) projections of input patterns of hand-written digits 0 and 8. (C) & (D): Higher principal-component (PC) projections of neuronal activities of a neuronal network corresponding to hand-written digits 0 and 8. The neuronal network was trained by synaptic competition.

**Appendix 1—figure 3.**
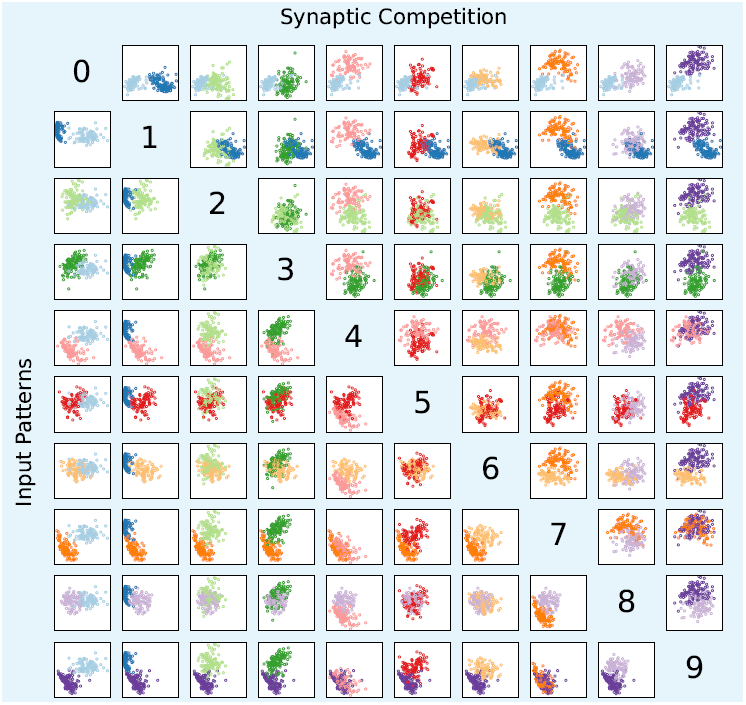
Comparisons of first and second principal components across digits. Lower triangle: principal components of input patterns. Upper triangle: principal components for activities of hidden units trained by synaptic competition. Horizontal axis: first principal components. Vertical axis: second principal components.

